# Multicancer analyses of short tandem repeat variations reveal shared gene regulatory mechanisms

**DOI:** 10.1101/2025.01.06.629343

**Authors:** Feifei Xia, Max Verbiest, Oxana Lundström, Tugce Bilgin Sonay, Michael Baudis, Maria Anisimova

**Author notes:** Contributing authors.

## Abstract

**Background:** Short tandem repeats (STRs) have been reported to influence gene expression across various human tissues. While STR variations are enriched in colorectal (CRC), stomach (STAD) and endometrial (UCEC) cancers, particularly in microsatellite instable (MSI) tumors, their functional effects and regulatory mechanisms on gene expression remain poorly understood across these cancer types.

**Results:** Here, we leverage whole-exome sequencing and gene expression data to identify STRs for which repeat lengths are associated with the expression of nearby genes (eSTRs) in CRC, STAD and UCEC tumors. Our analyses reveal that tumor STR profiles effectively capture both MSI phenotype and population structure. While most eSTRs are cancer-specific, shared eSTRs across multiple cancers exhibit consistent effects on gene expression. Notably, coding-region eSTRs identified in all three cancer types show positive correlations with nearby gene expression. We further validate the functional effects of eSTRs by demonstrating associations between somatic eSTR mutations and gene expression changes during the transition from normal to tumor tissues, suggesting their potential roles in tumorigenesis. Combined with DNA methylation data, we perform the first quantitative analysis of the interplay between STR variations and DNA methylation in tumors. We identify eSTRs where repeat lengths are associated with methylation levels of nearby CpG sites (meSTRs) and show that over 70% of eSTRs are significantly linked to local DNA methylation. Importantly, the effects of meSTRs on DNA methylation remain consistent across cancer types.

**Conclusions:** Overall, our findings enhance the understanding of how functional STR variations influence gene expression and DNA methylation. Our study highlights shared regulatory mechanisms of STRs across multiple cancers, offering a foundation for future research into their broader implications in tumor biology.

## Introduction

Short tandem repeats (STRs), also known as microsatellites, are repetitive sequences of 1 to 6 base pairs that are highly polymorphic and widely distributed throughout the human genome [1, 2]. They account for approximately 3% of the human genome [3]. STR mutation rates are orders of magnitude higher than existing estimations of the nucleotide substitution rates, thus representing a significant source of genetic variations [4–6]. Pathogenic expansions of STRs have been known to cause Mendelian disorders and neurological diseases, such as Fragile X syndrome and Huntington’s disease [7]. The underlying pathological mechanisms are diverse and depend on their genomic location, sequence composition and length [8, 9]. While coding-region STR expansions can directly lead to misfolding of the protein, the majority of studied STR expansions are in non-coding regions, potentially affecting gene expression through a variety of mechanisms, such as epigenetic silencing of the gene *FMR1* [10], acting as nucleosome positioning signals [11], altering the affinity of nearby DNA-binding sites [12]. Beyond Mendelian disorder, STRs have been proposed to contribute to complex traits in diverse organisms. Recently, multiple genome-wide association studies have revealed that STR length variations play a significant role in molecular and cellular processes, including gene expression [13–17], DNA methylation [18, 19], and alternative splicing [20].

STRs have been closely associated with cancer, particularly through their involvement in microsatellite instability (MSI) [21]. MSI arises from defects in the DNA mismatch repair (MMR) system, leading to an accumulation of mutations in STRs. MSIis a critical phenotype in multiple cancers, especially in colorectal, stomach, and endometrial cancer [22–24]. It has been detected in over 15% of patients in these cancers, and associated with favorable responses to immunotherapy [21]. STR variations in coding regions can result in frameshift mutations that may alter the function of tumor driver genes and also provide a substantial source of tumor-specific neoantigens [25, 26]. Given the prognostic and therapeutic significance of MSI, many studies have focused on accurate detection of MSI phenotype from next-generation sequencing and understanding the landscape of MSI across diverse cancer types [27–31]. However, despite the established roles of STRs in MSI detection and neoantigens generation, the regulatory functions of STR variations in influencing gene expression in cancers have not been fully characterized [32].

DNA methylation has emerged as another important diagnostic and prognostic marker for many cancers, including colorectal, stomach and endometrial cancer [33–35]. It has been demonstrated that DNA methylation plays a critical role in regulating gene expression by modifying transcription factor binding sites, acting either as a repressive or activating mark depending on the methylation region [36]. One of the key mechanisms leading to MSI is the promoter methylation of the DNA mismatch repair gene MLH1, which is an essential component of the mismatch repair system [37]. Furthermore, MSI tumors often overlap with the CpG Island Methylator Phenotype (CIMP), a distinct epigenetic subtype characterized by widespread CpG island hypermethylation [38]. Despite the well-established interactions, such as *MLH1* hypermethylation, and overlapping patterns between MSI and CIMP, the interplay between STR variations and local DNA methylation levels has not been quantitatively investigated in cancers.

Building on our previous identification of eSTRs (expression short tandem repeats) in CRC [32], we extended our analyses to stomach and endometrial cancers and incorporated DNA methylation data to explore the dual role of STRs functioning as eSTRs and meSTRs (methylation and expression short tandem repeats) across multiple cancers. Using tumor-derived STR genotypes, gene expression, and DNA methylation data, we performed quantitative trait locus (QTL) analyses to identify eSTRs and subsequently meSTRs where eSTR repeat lengths are associated with nearby DNA methylation levels. Our results revealed a strong concordance between eSTRs and meSTRs, with consistent regulatory effects across cancers. Overall, these findings highlight the multifaceted regulatory roles of STRs and suggest shared mechanisms underlying their influence on gene expression.

## Results

### STR profiles capture MSI phenotype and population structure in tumors

To explore the underlying patterns of STR length variations across CRC, STAD, and UCEC cancers, we utilized whole-exome sequencing (WES) data from 433, 439, and 464 paired normal and tumor samples obtained from The Cancer Genome Atlas (TCGA) database. We used GangSTR [39] to genotype STRs in each sample and then performed principal component analysis (PCA) on variable STRs for each cancer type to investigate the associations between STR length variations and molecular phenotypes (Fig. 1a). Across all three cancers, the first principal component (PC1) consistently separated samples based on MSI status, with MSI tumors clustering distinctly from microsatellite-stable (MSS) tumors. This separation underscores the strong relationship between MSI phenotype and STR length variations, particularly reflected in the percentage of STR somatic deletions (Fig. 1d). Notably, we observed a strong correlation between PC2 from tumor samples and PC1 from paired normal samples (Fig. 1b). This correlation validated the consistency of STR genotyping between tumor and normal tissues, confirming that tumor STR profiles represent authentic biological variation. Additionally, clear patterns of population structure were captured in both PC2 of tumor samples and PC1 of normal samples (Fig.1c, Additional file 1: Fig.S1), further supporting the reliability of the STR genotyping data from tumors. Besides the population structure, we also identified patterns related to technical artifacts (Additional file 1: Fig.S2), which were subsequently accounted for as covariates in the downstream association analyses.

**Fig. 1.**
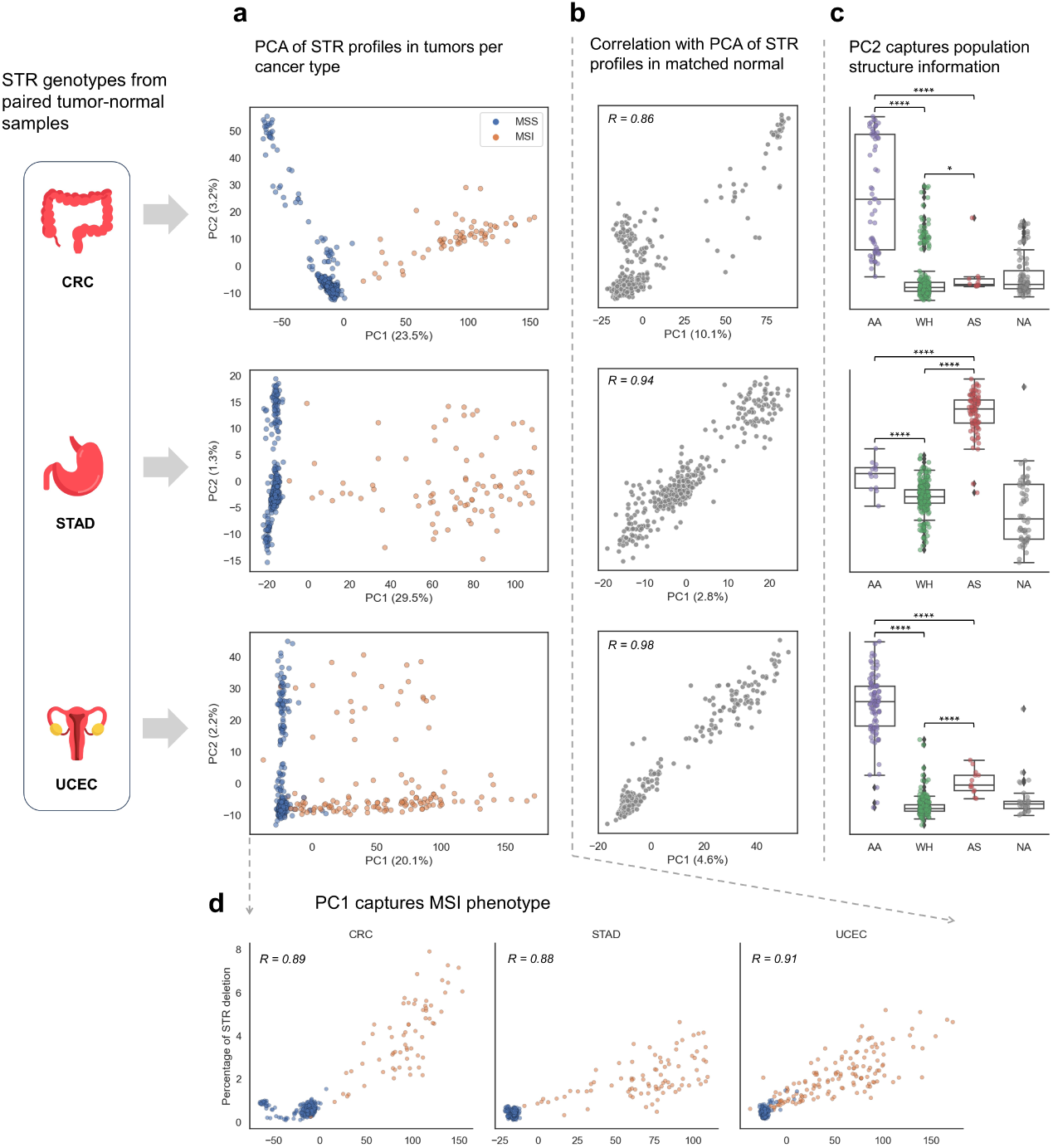
Tumor STR profiles capture MSI phenotype and population structure. **a** Principal component analysis plot of the first two principal components based on STR profiles from tumor samples per cancer type. The MSI status of each sample is indicated by colors. Numbers in brackets show the proportion of variance explained by each PC. **b** Pairwise comparisons of the PC2 from PCA of STR profiles in tumor samples and the PC1 from PCA of STR profiles in matched normal samples in three cancers, as shown by Pearson correlation coefficients (*P <* 1*e* − 10). **c** Boxplots showing the distribution of PC1 across population groups in each cancer type (AA: Black or African American; WH: white; AS: asian; NA: unknown). Two-sided Mann-Whitney-Wilcoxon tests were used to assess differences between population groups AA, WH and AS. Significant differences are indicated by asterisks (∗ : *p <* 0.05, ∗ ∗ ∗∗ : *p <* 1*e* − 4). **a,b and c** share the same y axes as shown in **a**. **d** Pairwise comparisons of the PC1 from PCA of STR profiles in tumor samples and the percentage of STR somatic deletion, as shown by Pearson correlation coefficients (*P <* 1*e* − 10).

### eSTRs identification and characterization

We performed an exome-wide analysis to identify associations between repeat length at each STR locus and the expression of nearby genes in CRC, STAD and UCEC tumors. Our analysis focused on 1326 tumor samples with highquality WES and RNA-sequencing data available from the TCGA database (Fig. 2a). After filtering low-quality STR calls (see Methods), we fitted linear models to test for associations between gene expression and the mean length of each STR located within the gene bodies or up to 5 kilobases (kb) upstream. The models were adjusted for gender, population structure, and technical covariates (see Methods). In total, 50283 STR-gene pair tests were performed across the three cancer types. After applying Benjamini–Hochberg multiple testing correction with a significance threshold of *FDR <* 0.05, we identified 1411, 382, and 108 significant STR-Gene associations in CRC, STAD, and UCEC respectively (Fig.2b, Additional file 2: Table S1). These STR loci were considered as eSTRs with putative regulatory roles. Over 60% of eSTRs were mononucleotide repeats located within intronic regions, with a small proportion located in coding or non-coding exons and untranslated regions (UTRs) (Fig.2c). These findings align with previous research showing that mononucleotide repeats are the most abundant STRs in the human genome [2] and represent a significant source of STR variability in MSI tumors [30, 40]. We next examined effect-size biases among STR-gene associations. Overall, we did not observe any significant bias in the direction of STR effects on gene expression. Among the eSTRs detected in CRC, we observed interesting associations. For instance, the repeat length of an intronic eSTR positively correlated with the expression level of UBR5 (Fig.2d), a gene that has been studied to be of clinical and biological significance in the progression of CRC [41, 42]. Similarly, we identified a negative association between the repeat length of an intronic eSTR and the expression of JAK2 (Fig.2e), which is involved in the pathogenesis of CRC [43, 44].

**Fig. 2.**
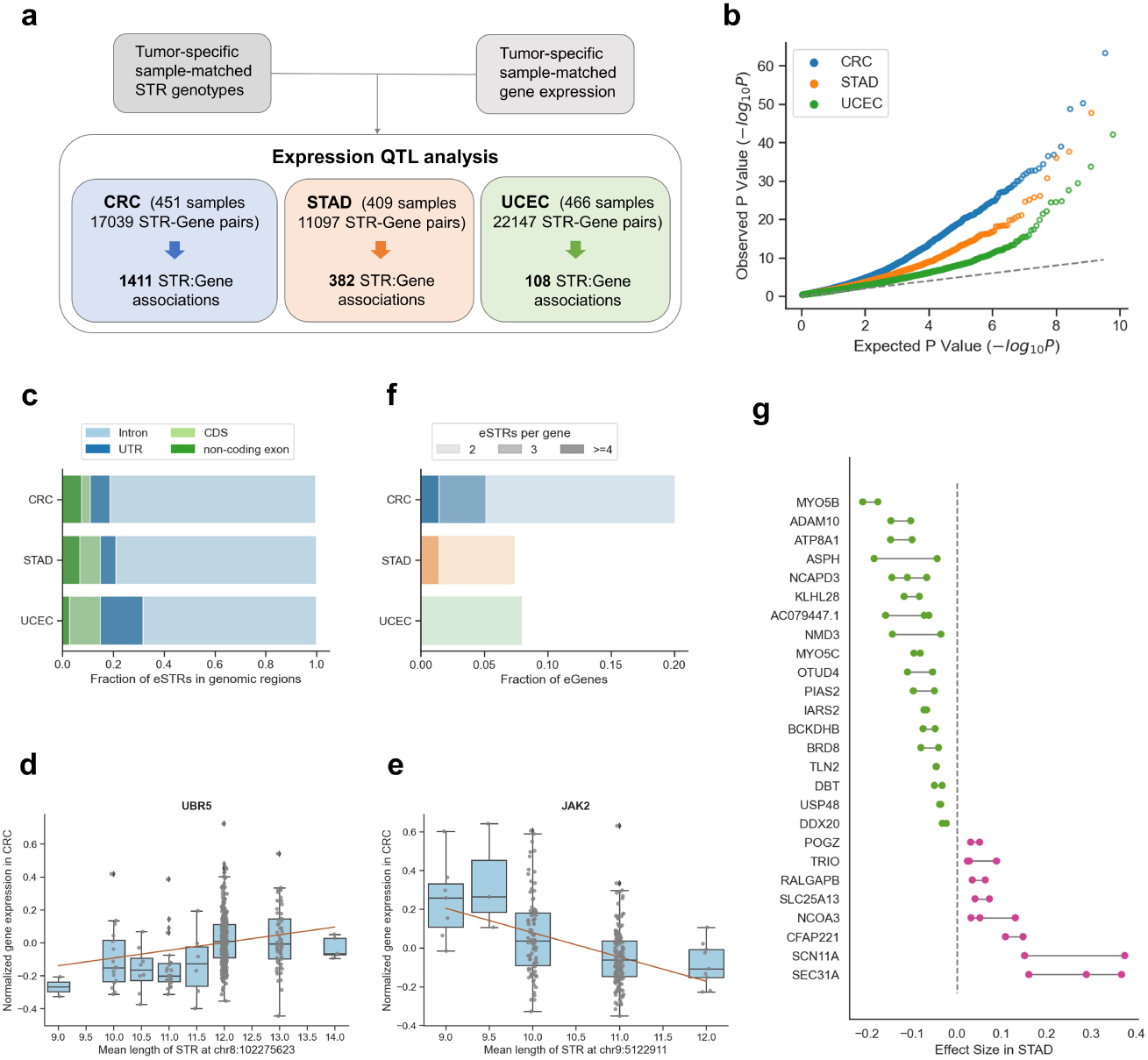
eSTR identification and characterization. **a** Schematic design of expression QTL analysis to identify eSTRs. We analyzed eSTRs using sample-matched gene expression and STR genotypes from whole exome sequencing data from TCGA CRC, STAD and UCEC. **b** eSTR association results. The quantile-quantile plot compares observed P values for each STR versus the expected uniform distribution for each tumor. **c** Barplots indicating fractions of eSTRs located within genomic regions (CDS, coding sequence; UTR, untranslated exon region) in CRC, STAD, UCEC tumors respectively. **d** Scatterplot showing an example of positive correlation between repeat length of eSTR in intronic region and UBR5 gene expression. **e** Scatterplot showing an example of negative correlation between repeat length of eSTR in intronic region and JAK2 gene expression. **f** Distribution of the number of eSTRs per gene, stratified by cancer type. Only genes with at least two identified eSTRs are shown in the plot. **g** Forest plot showing the consistent effects of multiple eSTRs on each gene. 26 genes with multiple eSTRs in STAD tumors are shown in the plot. Each dot represents an eSTR, with the direction of effects shown in pink (positive correlations) or green (negative correlations).

While most eGenes (genes associated with at least one eSTR) were linked to a single eSTR, there were 223, 26, 8 eGenes in CRC, STAD and UCEC associated with multiple eSTRs (Fig.2f). In CRC, some eGenes were associated with up to five eSTRs. Notably, these multiple eSTRs generally showed the same direction of correlation on gene expression. For example, 26 eGenes associated with multiple eSTRs showed concordant effect directions in STAD (Fig.2g). This pattern of consistent effects suggests that eSTRs are not simply tagging other causal variants, as indicated by previous studies [45]. However, exceptions in CRC included *ERAP2*, *KLKB1*, *TRAPPC4*, and *POLQ*, which were associated with eSTRs of different repeat unit lengths. This could be related to the hypothesis that distinct repeat classes may affect gene expression through different regulatory mechanisms [14]. Additionally, for eGenes *ST7*, *FAHD1*, and *WFDC3*, the associated eSTRs were assigned to multiple genes due to overlapping transcripts.

Compared to our previous study [32], which identified 1,295 significant STR-gene pairs, we confirmed 476 eSTRs here with our larger sample size and more robust confounder adjustments. While the previous study did not specifically investigate the consistent directional effects of multiple eSTRs on a single gene, this pattern of multiple eSTRs was also observed in the previous study except for gene *ATP6V1D*, *SYNE2* and *ERAP2*, reinforcing the reliability and biological relevance of this finding.

### Consistent effects of eSTRs on gene expression across cancers

Among the significant STR-gene associations, there were 1113, 351, 100 unique eGenes in CRC, STAD, UCEC respectively. To explore the regulatory roles of eSTRs on gene expression across cancers, we visualized the overlap of eGenes identified in each cancer type (Fig.3a). For each pair of cancers, we selected shared eGenes and computed the Pearson correlation between their effect sizes (Fig.3b). When eGenes were associated with multiple eSTRs, we averaged their effect sizes. Remarkably, 82.7%, 90% and 93.3% of overlapped eGenes were linked to at least one common eSTR. The effect sizes of these shared eSTRs and eGenes exhibited strong correlations across cancer types: CRC and STAD (*R* = 0.69*, P* = 2.26*e^−^*^20^), CRC and UCEC (*R* = 0.63*, P* = 1.8*e^−^*^4^), STAD and UCEC (*R* = 0.90*, P* = 5.01*e^−^*^6^). Our results showcase that certain eSTRs may regulate gene expression through shared regulatory mechanisms across different cancer types. This supports previous findings that most eSTRs act by global mechanisms across diverse human tissues [14].

**Fig. 3.**
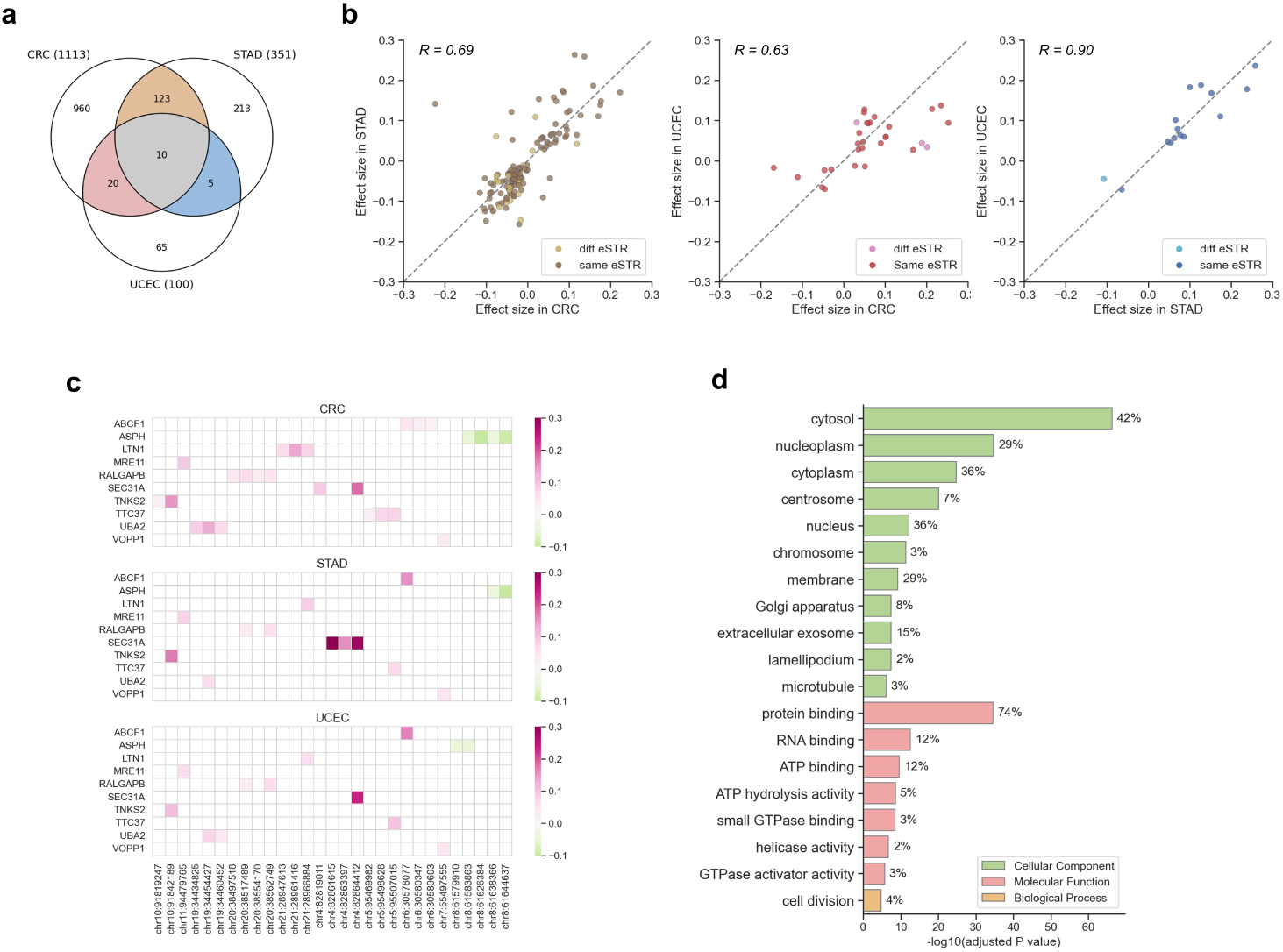
The effects of eSTRs on gene expression across cancers. **a** Venn diagram represents the overlap between eGenes in each cancer type. There are 1113, 351 and 100 unique eGenes detected in CRC, STAD and UCEC respectively. **b** Satterplots for comparison of effect sizes of overlapped eGenes between CRC and STAD, CRC and UCEC, STAD and UCEC (from left to right). Lighter dots (diff eSTR) denote the eGenes are not associated with common eSTR. Darker dots (same eSTR) denote the eGenes are associated with at least one common eSTR. **c** Heatmap showing the effect sizes of 10 eGenes that are detected in all three cancer types. Purple cells represent positive effects, green cells represent negative effects. **d** Enriched GO of 1113 eGenes identified in CRC from Gene Ontology enrichment analysis (*FDR <* 0.01).

Next, we examined the ten eGenes shared across all three cancers (Fig.3c). Interestingly, the expression of *ABCF1*, *LTN1*, *SEC31A*, and *TNKS2* positively correlated with eSTRs located in coding regions (Additional file 1: Fig.S5). STR variations in coding regions have been known to cause frameshift mutations, which can provide a substantial source for tumor-specific neoantigens [26, 46]. These findings suggest that eSTRs in coding regions may not only regulate gene expression but also contribute to the immunogenic landscape of cancer by facilitating the generation of tumor-specific neoantigens.

To further explore the biological processes and functions influenced by eSTRs, we performed Gene Ontology (GO) enrichment analysis on the eGenes identified in each cancer type (Additional file 3: Table S2). In CRC, the enrichment analysis revealed significant enrichment in the molecular function category *protein binding* (Fig.3d). In particular, 74%, 71%, and 77% of the eGenes detected in CRC, STAD, and UCEC respectively, were enriched under the *protein binding* category. These findings align with the previous observation [47, 48], which reported that genes containing variable repeats were frequently involved in protein binding. Moreover, eGenes from CRC and STAD showed significant enrichment in the *Phosphoprotein* and *Acetylation* categories, whereas eGenes from UCEC were exclusively enriched in the *Acetylation* category. These distinct enrichment patterns suggested a potential regulatory mechanism where eSTRs may influence gene expression through post-translational modifications (PTMs).

### eSTRs show higher mutability and regulate gene expression during tumorigenesis

We next sought to compare the somatic mutability of eSTRs and non-eSTRs using STR genotypes from paired samples. Due to the significant differences in STR mutability between MSI and MSS tumors, we assessed the mutability of eSTRs separately for MSI and MSS tumors. Similar to our previous study [32], we categorized the STRs based on their repeat unit size and allele length and then compared their mutability in each category. For example, in MSI tumors from CRC patients, eSTRs exhibited higher mutability than non-eSTRs in 25 of the total 31 categories. This was significantly higher than expected under a null distribution generated through random permutations of eSTR labels (Fig.4a, Additional file 1: Fig.S6). The trend of elevated mutability for eSTRs was consistent across nearly all MSI and MSS tumors, with the exception of MSS tumors in UCEC patients. Additionally, both in MSI and MSS patients, the mutability of eSTRs followed a descending trend from CRC, to STAD, and then to UCEC tumors.

**Fig. 4.**
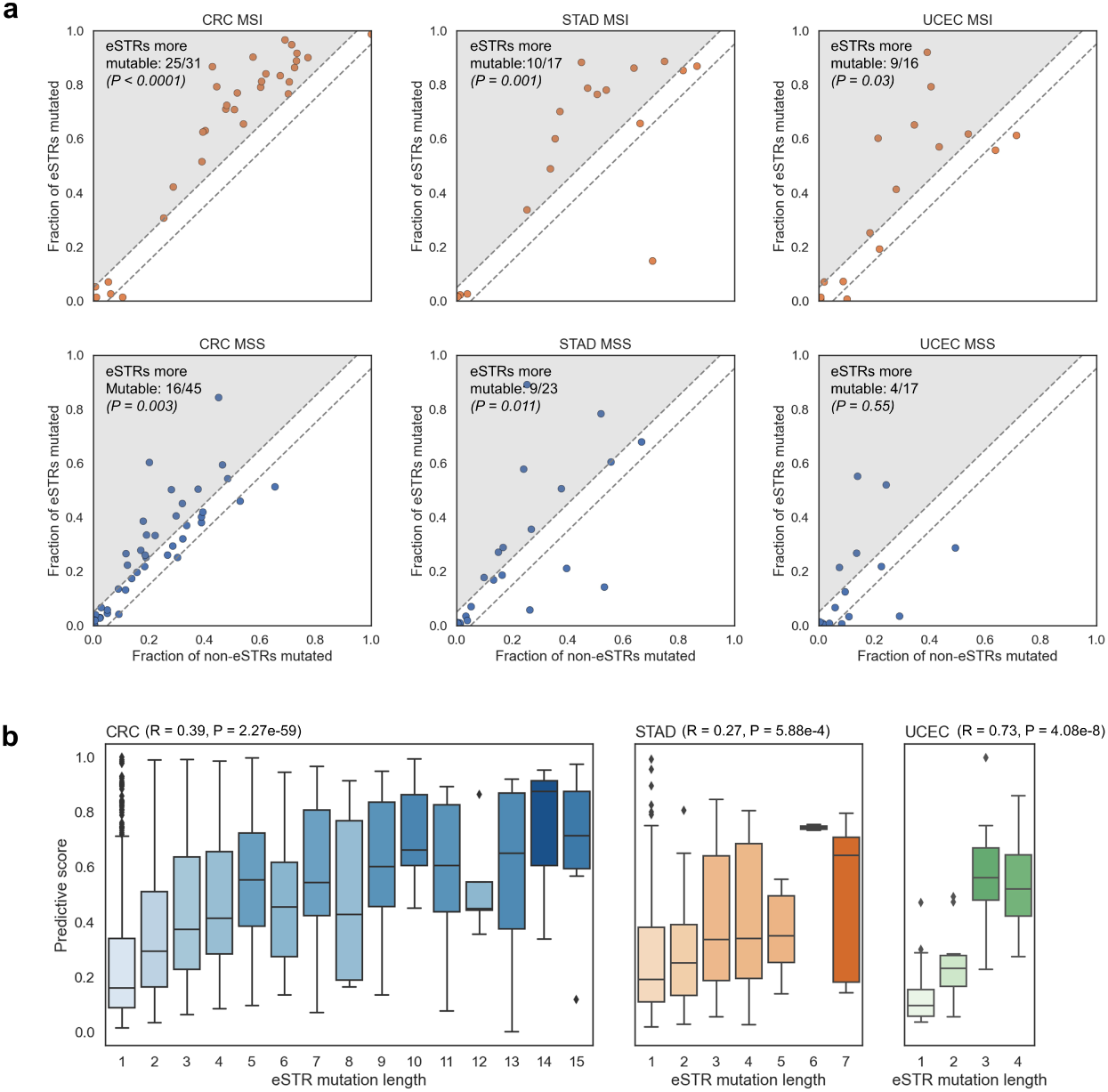
eSTRs have higher mutability and are predictive of gene expression change **a** Comparison of the mutability between eSTRs and non-eSTRs. The top row shows results for MSI patients and the bottom row shows results for MSS patients. Each dot in the scatterplot represents a STR category characterised by the STR unit size and allele length. The fraction of mutated non-eSTRs is shown on the x-axis, and the fraction of mutated eSTRs on the y-axis. Dots that fall between the dashed lines represent repeat types for which no difference in mutability between eSTRs and non-eSTRs was observed. For repeat types that fall in the shaded region, eSTRs were more mutable than non-eSTRs (their numbers are noted in the top left). P-values obtained from comparing the observed values to their respective null distributions using permutation tests are shown in the top left. **b** Boxplots showing the relationship between the eSTR mutation length and the predictive score of eSTR mutation prediction for gene expression change from normal to tumor samples. The x-axis represents different eSTR mutation lengths. Boxes are colored based on the prediction of gene expression change prediction. Larger mutation lengths lead to higher predictive score in predicting gene expression changes.

To further explore the functional consequences of eSTR mutability on gene expression, we focused on a subset of patients with matched gene expression data from both tumor and normal tissues. For each mutated eSTR, we calculated the eSTR mutation length (Δ*x*) and quantified the expression shift in the associated gene (Δ*y*) between the matched normal and tumor samples. We aimed to evaluate whether eSTR mutations could predict the direction and magnitude of corresponding gene expression changes during the transition from normal to tumor samples. The expected impact on gene expression changes was estimated by multiplying the eSTR mutation length (Δ*x*) with the adjusted effect sizes (*β^′^*) derived from our regression models. In general, the observed accuracy of predicting the direction of gene expression change was 55.6%, 57.5%, 53.3% for CRC, STAD, UCEC respectively, exceeding the random expectation of 50%. To assess the significance of these accuracies, we performed permutation tests for each cancer type by generating a null distribution from permutations of the eSTR effect sizes (*β^′^*). The resulting p-values (Additional file 1: Fig.S7) indicated that the observed accuracies were significantly higher than expected under the null hypothesis for CRC and STAD. Furthermore, we grouped the eSTR mutations by the absolute predicted impacts (|*β^′^* ∗ Δ*x*|) and calculated the average mutation impact for each group. We observed that eSTR mutations with higher average impacts resulted in higher accuracy in predicting the direction of gene expression changes (Additional file 1: Fig.S8a). For eSTR mutations with accurate direction predictions, we next evaluated their ability to predict the magnitude of corresponding gene expression changes. We calculated the predictive score (see Methods) for each eSTR mutation and demonstrated that the overall predictive scores were significantly higher than those of the baseline models (see Methods; Additional file 1: Fig.S9). When we again grouped them based on their absolute expected impacts, we observed a significantly increasing trend between the average mutation impacts and predictive scores across all cancer types (Additional file 1: Fig.S8b).

Finally, we wondered whether eSTR mutation length might affect their predictions about gene expression changes. We grouped the eSTR mutations based on their mutation lengths (Δ*x*) and tested for associations between eSTR mutation lengths and two metrics: the accuracy of predicting the direction of gene expression changes and the predictive score. Notably, we observed a significant positive correlation between eSTR mutation length and predictive score, but not with directional accuracy (Fig.4b; Additional file 1: Fig. S8c). Overall, our results further validated the linear relationship between eSTR length and gene expression, highlighting the functional role of eSTRs in mediating gene regulatory changes during tumorigenesis. For smaller eSTR mutation lengths, the effect on gene expression may be minimal or masked by other regulatory mechanisms. However, as the eSTR mutation length increased, the influence of eSTR mutation on gene expression became more pronounced.

### Further Evidence for a regulatory role of eSTR

MSI and MSS tumors exhibit distinct molecular features. MSI tumors are characterized by frequent point mutations and hypermethylation, whereas MSS tumors typically display high chromosomal instability [21, 23, 24, 49]. In addition to the variability of STRs, numerous other genetic and epigenetic factors may contribute to gene expression differences between MSS and MSI tumors. To provide further evidence of the regulatory role of eSTRs, we analyzed their effects in the context of the MSI phenotype. Specifically, we performed association analyses while conditioning the MSI phenotype of tumors for each cancer type. If the observed effects of eSTRs were solely attributable to the confounding influence of MSI, which can simultaneously affect both STR allele length and gene expression, then the conditioned effect of eSTRs should be randomly distributed compared to their unconditioned effects (Fig.5b). Alternatively, if the effects of eSTRs were independent of the MSI phenotype, the direction of conditioned effects should align with the unconditioned effects (Fig.5c). We considered eSTRs to be more likely causal regulators of gene expression if after conditioning, the direction of the associations stayed consistent and the associations remained statistically significant (*P <* 0.05). Using this approach, we found that 24.5%, 30.4% and 63.9% of eSTR signals were independent of MSI phenotype in CRC, STAD and UCEC respectively (Fig.5a, Additional file 4: Table S3).

**Fig. 5.**
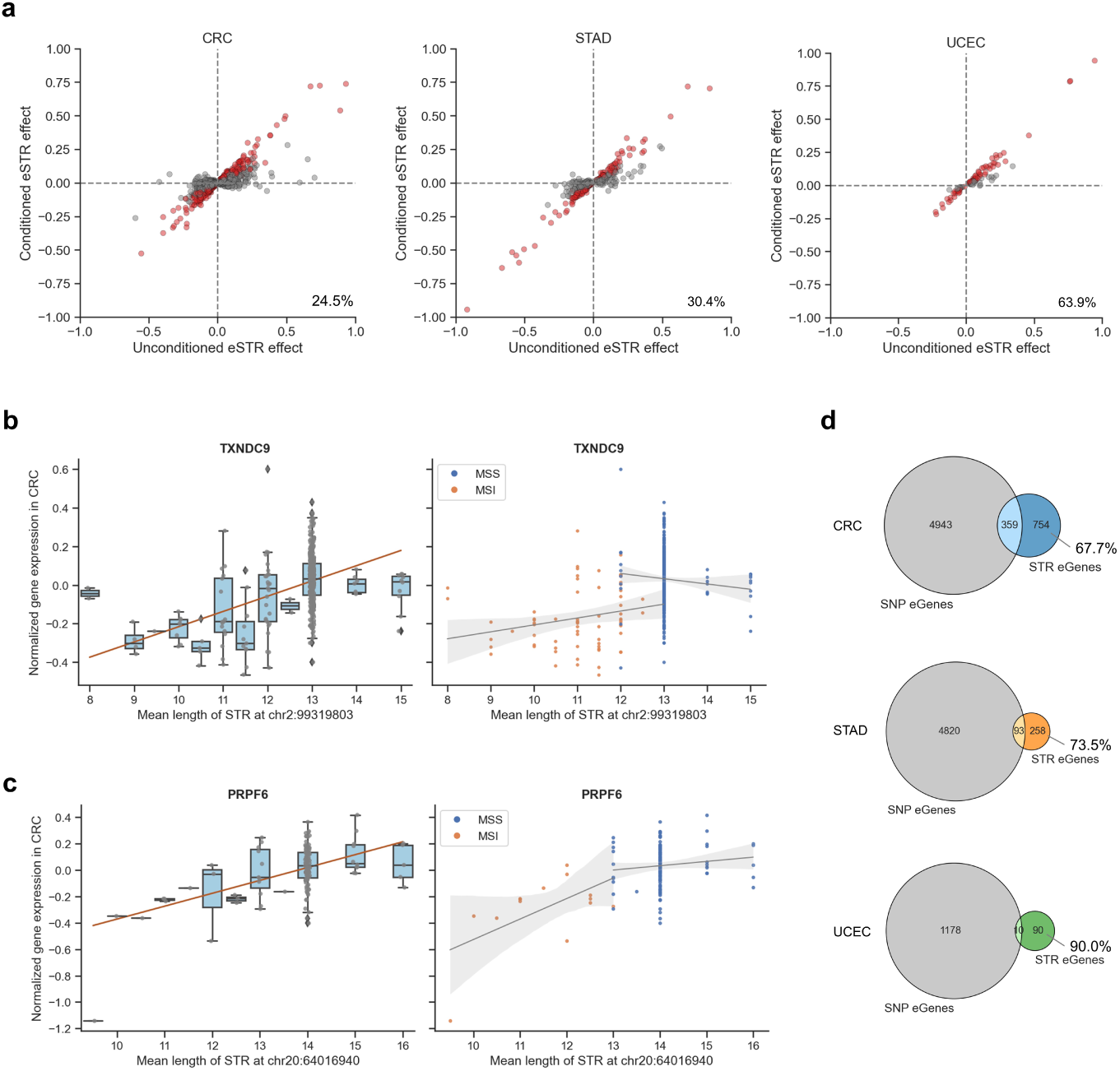
Further evidence for a regulatory role of eSTR. **a** Orignial eSTR effect versus conditioned eSTR effect. Red dots represent eSTRs whose directions of effect were consistent in both MSI tumors and MSS and whose associations remained significant (*P <* 0.05) upon conditioning for the MSI phenotype. Gray dots represent eSTRs that either became non-significant or showed opposite direction of effects after conditioning. **b** An example of STR:gene associations (TXNDC9), after conditioning on MSI phenotype, the direction of associations are different in MSS and MSI samples. **c** An example of STR:gene associations (PRPF6), after conditioning on MSI phenotype, the direction of associations remain the same and have statistical significance (*P <* 0.05). **d** Overlap of STR-associated eGenes and SNP-associated eGenes.

Previous studies have indicated that single nucleotide polymorphisms (SNPs) adjacent to mononucleotide repeats often show increased variability [50], thus the signals attributed to eSTRs might be explained by nearby SNPs in linkage disequilibrium [13, 14]. To evaluate this, we leveraged the database PancanQTL [51], which identified SNPs functioning as expression quantitative trait loci (eQTLs) across diverse cancer types. We examined the overlap between STR-associated eGenes and SNP-associated genes from the PancanQTL database in each cancer type. Our analysis revealed that the majority of eGenes, 67.7%, 73.5%, and 90.0% in CRC, STAD and UCEC respectively, did not overlap with SNP-associated genes (Fig.5d).

After filtering through both conditional and overlap analyses, 19.1%, 24.2% and 60.0% of the original eGenes in CRC, STAD and UCEC respectively, were retained. Notably, the genes *ABCF1*, *SEC31A*, *TNKS2* and *MRE11* still appeared in the overlap. Among these, *ABCF1*, *SEC31A* and *TNKS2* were associated with coding-region eSTRs. Collectively, these results suggest that eSTRs linked to these eGenes were more likely to as as independent casual regulators of gene expression.

### eSTRs are more likely to link to nearby DNA methylation

Functional STRs have previously been reported to influence nearby DNA methylation in the human genome [18, 19]. Additionally, the interactions between MSI and tumor DNA methylation have been studied in the pathogenesis of colorectal cancer [52]. Therefore, we sought to determine whether our eSTRs were enriched for associations with nearby DNA methylation. Using DNA methylation data from the TCGA database, we first mapped each STR locus to nearby CpG sites and then performed eSTR:CpG association analyses for each cancer type (see Methods). We identified that 84.9%, 81.3% and 72.6% of eSTRs were also meSTRs (eSTRs that are associated with nearby DNA methylation) in CRC, STAD and UCEC respectively (*FDR <* 0.05) (Additional file 5: Table S4), suggesting a strong concordance between eSTRs and meSTRs. To assess the enrichment of eSTRs for DNA methylation associations, we performed a comparative analysis with non-eSTRs. From the larger collection of non-eSTRs, we generated 1,000 background sets through random subsampling, ensuring each set matched the corresponding eSTR set in size. We then determined the percentage of mSTRs in which STR lengths were associated with DNA methylation levels of nearby CpG sites in these background sets, which served as null distributions. Notably, the proportion of meSTRs among eSTRs was 2–3 times higher than that observed in the null distributions (Fig. 6a), indicating a significant enrichment of DNA methylation associations for eSTRs.

**Fig. 6.**
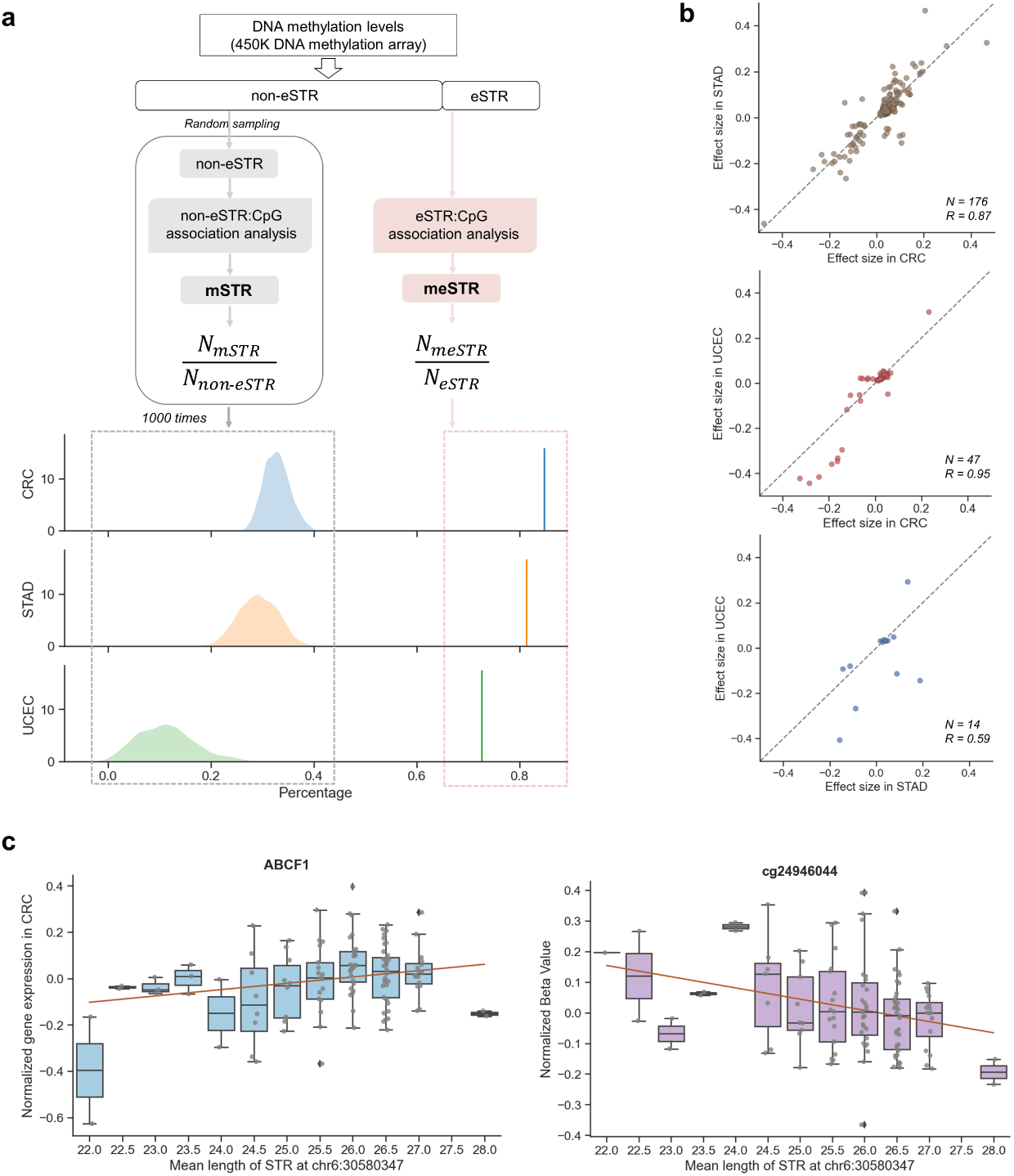
eSTRs are more likely to link to DNA methylation. **a** Illustration and comparison of mSTR/meSTR identification from non-eSTRs and eSTRs. To generate a null distribution, we randomly selected the amount of non-eSTRs as the same number of eSTRs per cancer type. The resulting distributions are shown in the left panel of the density plots. The proportions of meSTRs identified from eSTRs are shown in the right panel. **b** Comparison of effects of shared STR-CpG pairs between CRC and STAD, CRC and UCEC, STAD and UCEC (from left to right). Dots denote the same meSTR-CpG pairs between two cancers. **c** Scatterplots showing an example of the opposite correlation between the mean length of an eSTR with the gene ABCF1 expression (left plot) and nearby DNA methylation at CpG site cg24946044 (right plot).

Next, we investigated the functional characteristics of meSTRs to gain insight into their biological roles across the three cancer types. First, We examined effect-size biases in meSTR associations. Overall, meSTRs were more likely to show positive correlations between repeat length and DNA methylation in CRC and STAD (binomial one-sided *P <* 1.0*e^−^*^4^), but not in UCEC (binomial one-sided *P* = 0.92). Additionally, we observed that 77.7%, 82.4% and 59.7% of meSTRs in CRC, STAD and UCEC respectively displayed opposing correlation directions between meSTR:gene and meSTR:CpG associations. In most cases, increased repeat length of a meSTR was associated with lower gene expression and higher DNA methylation levels at one or more nearby CpG sites. These inverse correlations suggest that these meSTRs may regulate gene expression by modulating local DNA methylation. For example, the length of an intronic eSTR was positively correlated with the expression of gene *ABCF1* and negatively correlated with DNA methylation levels at a nearby CpG site (Fig.6c). Finally, we examined the consistency of meSTR associations across cancer types. The effects of overlapping meSTR associations were highly correlated between CRC and STAD (*R* = 0.87*, P* = 1.50*e^−^*^54^), CRC and UCEC (*R* = 0.95*, P* = 1.33*e^−^*^23^), STAD and UCEC (*R* = 0.59*, P* = 2.68*e^−^*^2^) (Fig.6b), similar to the relationship between overlapping eSTRs and gene expression. These consistent patterns further emphasized the robust effects of eSTRs on gene expression and highlighted potential shared regulatory mechanisms involving DNA methylation in these cancers.

## Methods

### Dataset collection

#### Whole exome sequencing data

Whole exome sequencing data (WES) for colon adenocarcinoma (COAD), rectum adenocarcinoma (READ), stomach adenocarcinoma (STAD), uterine corpus endometrial carcinoma (UCEC) from the TCGA database were accessed through dbGap under the study phs000178.v11.p8. The COAD and READ datasets were merged into a single colorectal cancer (CRC) dataset. According to the website GDAC (https://gdac.broadinstitute.org), FFPE (formalin fixed paraffin embedded) samples were not suitable for molecular analysis and were therefore excluded. WES data were downloaded using the GDC data transfer tool with default parameters. In total, there are 1110, 925 and 1118 samples available in CRC, STAD, UCEC cohorts respectively including tumor and normal samples. Batch information for the WES data, including sequencing plate, tumor purity, year, capture kit, and analyte type, was extracted from Search and retrieval in the GDC data portal. For duplicated samples from the same patient, the samples sequenced on the most recent plate were kept for downstream analyses.

#### Gene expression and DNA methylation

Gene expression (TPM normalized) and DNA methylation (Human methylation 450k) for CRC, STAD and UCEC cohorts were downloaded using the GDC data transfer tool from the Genomics Data Commons Data Portal (https://gdc-portal.nci.nih.gov/).

#### MSI phenotype

The MSI phenotypes (MSI-H, MSI-L/MSS) of CRC, STAD and UCEC tumors were obtained from the GDAC website (https://gdac.broadinstitute.org). Previous studies have suggested that MSI-L and MSS samples do not show distinct molecular and clinical features [53, 54]. Therefore, for simplicity, we merged MSI-L and MSS samples and collectively referred to them as MSS samples, while MSI-H samples were referred to as MSI.

### STR genotyping and preprocessing

STRs were genotyped from WES data using GangSTR [39] with the filtered STR panel [32] for the human protein-coding gene as the reference input. GangSTR was specially designed for short-read sequencing, which enabled the maximum likelihood estimation of repeat lengths beyond the read length based on various sources of information. Each sample was genotyped using GangSTR separately with the following additional parameters: --output-readinfo --nonuniform --include-ggl--verbose. Variant call format files (VCFs) were then filtered using DumpSTR [55] with the following settings: --gangstr-min-call-DP 20 --gangstr-max-call-DP 1000 --gangstr-filter-spanbound-only --gangstr-filter-badCI --zip--drop-filtered. The filtered VCFs were subsequently merged across all normal or tumor samples and processed for downstream analyses. The read lengths of WES data were 100/101 bp, 76 bp, 99/101 bp for CRC, STAD and UCEC cohorts respectively, which resulted in differences in the average number of genotyped STRs across cancer types. Due to relatively low mean coverage and low number of STR calls (*<* 10000 STR loci per sample), WES samples generated using the capture kit *Gapfiller 7m* were excluded from the CRC cohorts.

### Principal component analysis of STR profiles

For each cancer type, we performed principal component analysis (PCA) on a matrix **S***_n,m_* where *n* is the number of STR loci and *m* is the number of normal or tumor samples. Each element *c_i,j_* represents the mean STR allele length for a diploid call of the *j^th^* sample at the *i^th^* locus. *c_i,j_* is set to NaN when the *i^th^* STR call of the *j^th^* sample is missing. Before further analysis, STR loci missing in more than 50% of the total *m* samples were discarded. Additionally, STR loci with length variation ¡ 0.1 across *m* samples were also excluded. The filtered matrices were then imputed using k-nearest neighbors with the parameters n neighbors=5, weights=distance, implemented in the KNNImputer function from the Python scikit-learn library [56]. The imputed matrices were subsequently used as input for PCA analyses using the Python scikit-learn library.

### STR somatic mutation calling

Paired tumor and normal samples from the same patients were used for STR somatic mutation calling. There were 433 (MSS: 363; MSI: 70), 439 (MSS: 354; MSI: 85), and 464 (MSS: 323; MSI: 141) available paired samples for CRC, STAD, UCEC respectively. The biallelic STR genotypes at each locus were compared between paired normal and tumor samples. We separately analyzed deleted and inserted STR loci, where at least one allele in the tumor sample was shorter or longer than in the paired normal sample. Consistent with our previous findings, the elevated STR mutation percentages observed in MSI tumors were solely driven by deletions in all three cancer types (Additional file 1: Fig.S3). Similarly, we removed STR loci where the tumor sample appeared homozygous for an allele but not present in its matched normal sample. These calls could arise from heterozygous loci that were spuriously called as homozygous due to allele dropout [57].

### eSTRs identification

For each cancer type, we filtered out STR genotypes called in fewer than 10% of total tumor samples and then restricted our analyses to variable STRs with a standard deviation of repeat lengths *>* 0.1 within the cohort. We mapped STRs to a 5kb upstream window of gene regions to define STR-gene pairs. We excluded genes with median TPM values of zero and the remaining expression values were quantile normalized separately for each cancer type to a standard normal distribution. Analyses were further restricted to protein-coding genes based on GENCODE v.36 annotation. Altogether, 17039, 11097 and 22147 STR-gene pairs remained in 451 CRC, 409 STAD and 466 UCEC tumors respectively. For each STR-gene pair, we fitted a linear regression model between STR mean allele length and gene expression levels:

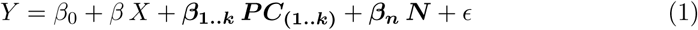

where *X* denotes STR genotypes, specifically mean allele length at the STR locus, *Y* denotes gene expression levels, *β* denotes the effect size and *ɛ* is the error term. To control for technical variations in expression, we applied PCA correction for gene expression [58]. To ascertain the number of principal components (PCs) that capture technical variability, we carried out multiple rounds of association analyses for each cancer type separately, each time incorporating 0,1,2,3,4,5 PCs as covariates. This approach identified 2, 0, and 2 PCs for CRC, STAD, and UCEC respectively, as optimal covariates for maximizing eSTR discovery (Additional file 1: Fig.S4). **N** represents a vector of additional covariates, including *population structure, gender* and *year*. To correct for *population structure*, we used inferred population stratification from an aggregated model as covariates for patients without self-reported population [59]. *Year* represents technical variation identified in STR genotypes (Additional file 1: Fig.S2). A separate regression analysis was performed for each STR-gene pair in each tumor type. Linear regression was fitted using the Ordinary Least Squares (OLS) function from the Python statsmodels.api module [60]. Samples with missing STR genotypes or expression values were excluded from each regression analysis. To reduce the effect of outlier STR genotypes, we removed samples with genotypes observed in fewer than three samples. To control the false discovery rate (FDR), we applied Bonferroni multiple testing correction for the number of evaluated STR-gene pairs tested within each cancer type. eSTRs were defined as STR loci where the mean allele length was significantly associated with the expression level of a nearby gene (*FDR <* 0.05).

### Cross-cancer eSTR analysis

To compare the effects of eSTRs across different cancer types, we first identified overlapping eGenes that showed significant associations with eSTRs in multiple cancers. We then applied regression analysis to calculate residual gene expression by accounting for the aforementioned covariates. Using these residuals, we estimated the posterior effect sizes of each eSTR in each cancer type. Subsequently, we performed pairwise correlation analyses of the adjusted effect sizes for the overlapping eGenes across cancer types. For eGenes associated with multiple eSTRs, we averaged the effect sizes to ensure a consistent and comparable evaluation.

### GO enrichment analysis

To gain insight into the biological functions and pathways associated with the identified eGenes, we performed Gene Ontology (GO) enrichment analysis using DAVID (Database for Annotation, Visualization, and Integrated Discovery) [61]. The analysis was conducted separately for each cancer type. We focused on GO categories for biological process (BP), molecular function (MF), and cellular component (CC) to identify overrepresented functional terms. GO terms with FDR corrected p-values *<* 0.05 were considered statistically significant in CRC. However, due to the smaller number of eGenes from STAD and UCEC tumors, no significantly enriched GO terms were identified for these cancer types.

### eSTR mutability analysis

For the STR loci included in the association analyses, we compared the mutability of eSTRs and non-eSTRs separately in MSI and MSS tumors. Following our previous approaches [32], we grouped all the STRs based on their unit size and reference allele length. For each repeat type, we retrieved all mutations of eSTRs and non-eSTRs for which somatic mutations had been previously calculated. To ensure robust analysis, repeat types in which STRs were observed fewer than 50 times were excluded for each cancer type. We then calculated the fraction of mutated eSTRs and non-eSTRs within each repeat type. We defined eSTRs as more mutable within a given repeat type when the fraction of mutated eSTRs exceeded the fraction of mutated non-eSTRs by at least 0.05. To test whether the observed number of repeat types with increased eSTRs mutability was significantly different from random expectation, we repeated the analysis for 10000 permutations. In each permutation, the labels indicating which loci were eSTRs were randomly shuffled, and the fraction of repeat types with an increased eSTRs mutability was recorded. This process generated null distributions, which served as baseline models for comparison. The observed fraction of repeat types with increased mutability in eSTRs was then compared to the null distributions using permutation tests to determine statistical significance.

### eSTR somatic mutation and gene expression change

Matched gene expression data were available in 33, 33, 19 paired tumor and normal samples for CRC, STAD and UCEC cohorts. Among these paired samples, we detected 3285, 302 and 94 somatic mutations in putative eSTRs using the STR mutation calling method described above. For each eSTR and its corresponding eGene, gene expression changes were determined by calculating the differences between paired tumor and normal samples after performing quantile normalization separately in the normal and tumor samples. Using the eSTR mutation lengths Δ*x* and the corresponding expression change Δ*y* of the associated eGenes, we assessed whether our linear models could predict gene expression changes in response to somatic eSTR mutations both directionally and quantitatively. To evaluate directional predictions, we determined whether the actual direction of gene expression change Δ*y* matched the predicted change based on the eSTR mutation length Δ*x* and the adjust coefficient *β^′^*. We recalculated the coefficients *β^′^* for each eSTR based on the residuals of adjusted gene expression values *Y ^′^*. A directional prediction was considered correct if both the gene expression change (Δ*y*) and the predicted change (*β^′^* ∗ Δ*x*) showed the same sign (both positive or negative). For eSTR mutations that correctly predicted the direction, we further calculated the predictive score to assess the accuracy of quantitatively predicting gene expression changes, as defined by the formula:

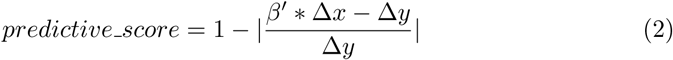

Negative predictive scores were observed for 11.5%, 10.6%, and 11.7% of eSTR mutations in CRC, STAD, and UCEC. These negative values likely reflect noise or other factors influencing gene expression. We retained only the eSTR mutations with positive predictive scores for visualization.

### Conditional analysis

Given that the MSI phenotype can affect both STR length variation and gene expression, we incorporated MSI status as a covariate in our regression models to account for its confounding effect. Specifically, we conducted a conditional analysis to determine whether the association between STR length variation and gene expression is independent of MSI phenotype. For each eSTR, we performed the following analysis:

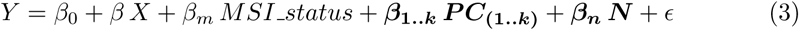

*MSI Status* is a binary variable indicating the microsatellite instability phenotype. eSTRs were considered as regulators of gene expression independent of MSI phenotype when the resulting association upon conditioning was still significant (nominal *P <* 0.05) and had the same directionality as in the unconditioned analysis.

### Overlap of eGenes and SNP-associated genes

SNP-associated genes in COAD, READ, STAD, and UCEC were downloaded from the database PancanQTL [51]. The database conducted expression quantitative trait locus (eQTL) analysis using gene expression data and SNP genotypes, and comprehensively provided the results of eQTLs from the TCGA. We filtered the SNP-gene association with a threshold of *FRD <* 0.05.

### meSTRs identification and analysis

Among the tumors we analyzed, DNA methylation data was available for 393, 395 and 431 samples in CRC, STAD and UCEC respectively. The methylation levels were represented by *β*-values ranging from zero (unmethylated) to one (fully methylated). To ensure the reliability of the methylation measurements, we excluded any probe that mapped to multiple genomic locations as described by [62]. Annotations for Illumina’s 450k methylation arrays were downloaded using R [63] and the minfi package [64]. The annotations were lifted to the hg38 reference genome using bedtools [65]. CpG sites with a minimum call rate below 80% were filtered out. For this analysis, we selected CpG sites located within a 5 kb upstream region or gene regions corresponding to each eSTRs. We performed association testing for 33910, 10672, and 2598 eSTR:CpG pairs in CRC, STAD and UCEC, controlling for gender, population structure and technical variations (as described earlier). Multiple testing corrections for the number of eSTR:CpG pairs evaluated were applied using the Benjamini-Hochberg method.

To assess whether eSTRs are more likely to be linked to DNA methylation, we generated background sets separately from 11879, 8223, 16832 non-eSTRs in CRC, STAD and UCEC. For each background set, we randomly sampled 1000 subsets of non-eSTRs with replacement, ensuring the number of non-eSTRs in each subset matched the number of corresponding eSTRs in each cancer type. We then mapped non-eSTR:CpG pairs using the same procedure applied for eSTRs and conducted identical association analyses. Subsequently, We quantified the proportion of mSTRs within each subset of non-eSTRs. The proportions of mSTRs within the background set for each cancer type established a baseline distribution, representing expectations under the null hypothesis.

To compare the effects of meSTRs across different cancer types, we first selected the eSTR:CpG pairs showing significant associations in multiple cancer types. We then applied linear regression analysis to calculate the residual methylation levels, accounting for relevant covariates. Using these residuals, we estimated the posterior effect sizes for each overlapped pair. Subsequently, we performed pairwise correlation analyses between the effect-size estimates for the overlapping eSTR:CpG pairs across cancer types to examine consistency in meSTR effects.

## Discussion

This study provides a comprehensive analysis of STR variations and their functional roles in CRC, STAD, and UCEC tumors. To our knowledge, this is the first study to analyze STRs across three distinct cancer types while integrating gene expression and DNA methylation data. This multi-cancer, multi-omics approach allowed us to uncover not only the functional impact of eSTRs on gene expression and DNA methylation, but also highlighted shared regulatory mechanisms through eSTRs and meSTRs conserved across cancers. These overlapping eSTRs likely represent a core set of key regulatory elements across cancers.

Our integrated analysis provided valuable insights into the regulatory roles of STRs. We first revealed that tumor STR profiles capture MSI phenotype and population structure, validating the robustness of STR genotyping in tumor samples. Next, we performed association analyses between STR repeat length and nearby gene expression and identified 1383, 370, and 107 unique eSTRs in CRC, STAD, and UCEC tumors respectively. We then used conditional analyses accounting for MSI phenotype and overlapping analyses with SNP-associated genes to investigate the causality of our putative eSTRs, which determined that 19.1%, 24.2%, 60.0% of them are more likely to act as independent regulators of nearby gene expression. We further demonstrated that eSTRs show higher mutability and their mutation lengths are strongly associated with gene expression changes during the transition from normal to tumor tissues, particularly those with larger mutation lengths. These findings provided additional evidence for the regulatory role of eSTRs and indicated that eSTR mutations may play a crucial role in driving gene expression alterations during tumorigenesis.

Our findings have several important implications for understanding the regulatory mechanisms of STRs on gene expression. First, by integrating DNA methylation data, we demonstrated that over 70% eSTRs were significantly associated with methylation levels of nearby CpG sites, forming what we termed meSTRs. Notably, the regulatory effects of overlapping eSTRs on gene expression were consistent across cancers. Similarly, overlapping meSTRs exhibited strong concordance in their influence on DNA methylation. These consistent patterns across cancers suggest that eSTRs may influence gene expression through epigenetic modulation in tumors, specifically by altering the methylation levels of adjacent CpG sites. Second, functional enrichment analyses of eGenes showed significant enrichment in post-translation modifications (PTMs). They are critical for modulating protein-protein interactions, protein stability, and localization in cancers [66]. The observed enrichment suggests that eSTRs may regulate gene expression by affecting PTM-related processes, such as transcriptional activation or repression, chromatin accessibility, and protein-protein interactions. While we proposed these two primary mechanisms—transcription factor modulation and methylation changes, as pathways through which eSTRs influence gene expression, our findings do not exclude the possibility of other regulatory mechanisms or complex interactions between these mechanisms. Future studies integrating additional multiomics data could help unravel these interactions and clarify the full spectrum of eSTR regulatory functions in cancer.

Our study also revealed a dual role for coding-region eSTRs identified in all the three cancers. We observed that the expression of genes *ABCF1*, *LTN1*, *SEC31A*, and *TNKS2* was positively correlated with the allele length of eSTRs located in the coding regions. Among these genes, *SEC31A* has been identified as one of five genes encoding shared immunogenic frameshift mutations in colorectal, stomach and endometrial MSI tumors [67]. Additionally, *LTN1* was among the six candidate genes that were predicted and validated to generate shared frameshift peptides with functionally confirmed immunogenicity in both colorectal and stomach MSI tumors [68]. Furthermore, *TGFBR2* and *SLC35F5*, which showed positive associations with eSTRs located in coding regions in both colorectal and stomach cancer, were also among these six candidate genes. Collectively, we hypothesize that eSTRs in coding regions may have a dual impact: (1) affecting gene expression, and (2) generating tumor-specific neoantigens through frameshift mutations, particularly in MSI tumors. This dual role could have further diagnostic and therapeutic implications for understanding the functional significance of STRs in cancer biology, offering insights into immunotherapy treatment for multiple cancers.

There are several limitations in our study. (1) The datasets we used for STR genotyping were derived from whole-exome sequencing, which is insufficient to capture all STRs in the genome, particularly those located in non-coding regions. As a result, certain potentially functional STRs may have been missed, limiting the scope of our analysis. Future studies employing whole-genome or long-read sequencing would allow for a more comprehensive exploration of STRs and their regulatory roles across cancers. (2) Although we performed conditional analyses and investigated overlap with SNP-associated genes to strengthen the evidence for identified eSTR signals, these analyses are susceptible to false negatives due to the stringent conditions used. (3) While our analyses were based on the assumption of linear relationships between STR length variations and gene expression, it is possible that the relationship is non-linear or follows alternative patterns that our models did not account for. Exploring non-linear associations in future research could reveal additional layers of regulatory complexity. (4) We demonstrated that eSTRs are more likely to influence the methylation levels of nearby CpG sites. However, the complex relationships among STR variations, DNA methylation and gene expression remain unclear. Further studies incorporating multi-omics data and advanced models are needed to unravel the mechanisms underlying their interactions.

## Conclusions

Overall, our integrated analyses of STR variations, gene expression and DNA methylation provide valuable insights into the functional role of STRs among colorectal, stomach and endometrial cancers. Our findings suggest that STRs may influence gene expression and DNA methylation through shared regulatory mechanisms in these cancers, highlighting their potential significance as key regulators in tumorigenesis. These results lay the groundwork for future studies to explore the clinical relevance of STRs as biomarkers or therapeutic targets.

## Data availability

The analyses were primarily performed using Python 3.10.4. Scripts for analyses and reproducing figures are available here: https://github.com/acg-team/multicancer_STR. Data from the COAD, READ, STAD, UCEC cohorts were downloaded from the GDC knowledge base (data release version 38.0). Access to restricted TCGA data was granted under dbGaP study phs000178.v11.p8. Summary statistics of STRs in the study will be available in the database WebSTR [69].

## Supplementary information

Additional file 1: Figure S1-S9.

Additional file 2: Table S1, eSTRs in CRC, STAD and UCEC;

Additional file 3: Table S2, enrichment analysis of eGenes

Additional file 4: Table S3, conditional analysis

Additional file 5: Table S4, meSTRs in CRC, STAD and UCEC

## Acknowledgements

FX, MAV, OL were supported by SNSF Sinergia grant CRSII5 193832 to MA. This work was also supported by the EU Horizon 2020 research and innovation program under the Marie Skl-odowska-Curie grant agreement No. 823886.

## Declarations

The authors declare no competing interests.

## Notes

### Competing Interest Statement

The authors have declared no competing interest.

## References

[1] Ellegren, H. Microsatellites: simple sequences with complex evolution. Nature Reviews Genetics 5, 435–445 (2004). URL https://www.nature.com/articles/nrg1348. Number: 6 Publisher: Nature Publishing Group.

[2] G, T., Z, G. & J, J. Microsatellites in different eukaryotic genomes: survey and analysis. Genome research 10 (2000). URL https://pubmed.ncbi.nlm.nih.gov/10899146/. Publisher: Genome Res.

[3] Fan, H. & Chu, J.-Y. A Brief Review of Short Tandem Repeat Mutation. Genomics, Proteomics & Bioinformatics 5, 7 (2007). URL https://www.ncbi.nlm.nih.gov/pmc/articles/PMC5054066/. Publisher: Oxford University Press.

[4] Sun, J. X., et al. A direct characterization of human mutation based on microsatellites. Nature Genetics 44, 1161–1165 (2012). URL https://www.nature.com/articles/ng.2398. Number: 10 Publisher: Nature Publishing Group.

[5] Willems, T., et al. The landscape of human STR variation. Genome Research 24, 1894 (2014). URL https://www.ncbi.nlm.nih.gov/pmc/articles/PMC4216929/. Publisher: Cold Spring Harbor Laboratory Press.

[6] Redelings, B. D., Holmes, I., Lunter, G., Pupko, T. & Anisimova, M. Insertions and Deletions: Computational Methods, Evolutionary Dynamics, and Biological Applications. Molecular Biology and Evolution 41 (2024). URL 10.1093/molbev/msae177. Publisher: Oxford Academic.

[7] Mirkin, S. M. Expandable DNA repeats and human disease. Nature 447, 932–940 (2007). URL https://www.nature.com/articles/nature05977. Number: 7147 Publisher: Nature Publishing Group.

[8] Gymrek, M. A genomic view of short tandem repeats. Current Opinion in Genetics & Development 44, 9–16 (2017). URL https://linkinghub.elsevier.com/retrieve/pii/S0959437X16301538.

[9] Uguen, K., Michaud, J. L. & Génin, E. Short Tandem Repeats in the era of next-generation sequencing: from historical loci to population databases. European Journal of Human Genetics 32, 1037–1044 (2024). URL https://www.nature.com/articles/s41431-024-01666-z. Number: 9 Publisher: Nature Publishing Group.

[10] Xs, L., et al. Rescue of Fragile X Syndrome Neurons by DNA Methylation Editing of the FMR1 Gene. Cell 172 (2018). URL https://pubmed.ncbi.nlm.nih.gov/29456084/. Publisher: Cell.

[11] T, R.-S., et al. Manipulating nucleosome disfavoring sequences allows fine-tune regulation of gene expression in yeast. Nature genetics 44 (2012). URL https://pubmed.ncbi.nlm.nih.gov/22634752/. Publisher: Nat Genet.

[12] Afek, A., Schipper, J. L., Horton, J., Gordân, R. & Lukatsky, D. B. From the Cover: ProteinDNA binding in the absence of specific base-pair recognition. Proceedings of the National Academy of Sciences of the United States of America 111, 17140 (2014). URL https://www.ncbi.nlm.nih.gov/pmc/articles/PMC4260554/. Publisher: National Academy of Sciences.

[13] Gymrek, M., et al. Abundant contribution of short tandem repeats to gene expression variation in humans. Nature Genetics 48, 22–29 (2016). URL https://www.nature.com/articles/ng.3461. Number: 1 Publisher: Nature Publishing Group.

[14] Fotsing, S. F., et al. The impact of short tandem repeat variation on gene expression. Nature Genetics 51, 1652–1659 (2019). URL https://www.nature.com/articles/s41588-019-0521-9. Number: 11 Publisher: Nature Publishing Group.

[15] Jakubosky, D., et al. Properties of structural variants and short tandem repeats associated with gene expression and complex traits. Nature Communications 11, 2927 (2020). URL https://www.nature.com/articles/s41467-020-16482-4. Number: 1 Publisher: Nature Publishing Group.

[16] Shi, Y., et al. Characterization of genome-wide STR variation in 6487 human genomes. Nature Communications 14, 1–18 (2023). URL https://www.nature.com/articles/s41467-023-37690-8. Number: 1 Publisher: Nature Publishing Group.

[17] Ca, H., et al. Short tandem repeats bind transcription factors to tune eukaryotic gene expression. Science (New York, N.Y.) 381 (2023). URL https://pubmed.ncbi.nlm.nih.gov/37733848/. Publisher: Science.

[18] Quilez, J., et al. Polymorphic tandem repeats within gene promoters act as modifiers of gene expression and DNA methylation in humans. Nucleic Acids Research 44, 3750 (2016). URL https://www.ncbi.nlm.nih.gov/pmc/articles/PMC4857002/. Publisher: Oxford University Press.

[19] Martin-Trujillo, A., Garg, P., Patel, N., Jadhav, B. & Sharp, A. J. Genome-wide evaluation of the effect of short tandem repeat variation on local DNA methylation. Genome Research 33, 184–196 (2023). URL https://genome.cshlp.org/content/33/2/184. Company: Cold Spring Harbor Laboratory Press Distributor: Cold Spring Harbor Laboratory Press Institution: Cold Spring Harbor Laboratory Press Label: Cold Spring Harbor Laboratory Press Publisher: Cold Spring Harbor Lab.

[20] Hamanaka, K., et al. Genome-wide identification of tandem repeats associated with splicing variation across 49 tissues in humans. Genome Research 33, 435–447 (2023). URL https://genome.cshlp.org/content/33/3/435. Company: Cold Spring Harbor Laboratory Press Distributor: Cold Spring Harbor Laboratory Press Institution: Cold Spring Harbor Laboratory Press Label: Cold Spring Harbor Laboratory Press Publisher: Cold Spring Harbor Lab.

[21] Li, K., Luo, H., Huang, L., Luo, H. & Zhu, X. Microsatellite instability: a review of what the oncologist should know. Cancer Cell International 20, 16 (2020). URL 10.1186/s12935-019-1091-8.

[22] Boland, C. R. & Goel, A. Microsatellite Instability in Colorectal Cancer. Gastroenterology 138, 2073 (2010). URL https://www.ncbi.nlm.nih.gov/pmc/articles/PMC3037515/. Publisher: NIH Public Access.

[23] Puliga, E., Corso, S., Pietrantonio, F. & Giordano, S. Microsatellite instability in Gastric Cancer: Between lights and shadows. Cancer Treatment Reviews 95 (2021). URL https://www.cancertreatmentreviews.com/article/S0305-7372(21)00023-2/fulltext. Publisher: Elsevier.

[24] Kanopiene, D. et al. Endometrial cancer and microsatellite instability status. Open Medicine 10, 70–76 (2014). URL https://www.ncbi.nlm.nih.gov/pmc/articles/PMC5152958/.

[25] Woerner, S. M., et al. SelTarbase, a database of human mononucleotide-microsatellite mutations and their potential impact to tumorigenesis and immunology. Nucleic Acids Research 38, D682–D689 (2010). URL https://www.ncbi.nlm.nih.gov/pmc/articles/PMC2808963/.

[26] Turajlic, S., et al. Insertion-and-deletion-derived tumour-specific neoantigens and the immunogenic phenotype: a pan-cancer analysis. The Lancet Oncology 18, 1009–1021 (2017). URL https://www.thelancet.com/journals/lanonc/article/PIIS1470-2045(17)30516-8/fulltext. Publisher: Elsevier.

[27] Kim, T.-M., Laird, P. W. & Park, P. J. The Landscape of Microsatellite Instability in Colorectal and Endometrial Cancer Genomes. Cell 155, 858–868 (2013). URL https://www.sciencedirect.com/science/article/pii/S0092867413012919.

[28] Bonneville, R., et al. Landscape of Microsatellite Instability Across 39 Cancer Types. JCO Precision Oncology 1–15 (2017). URL https://ascopubs.org/doi/full/10.1200/PO.17.00073. Publisher: Wolters Kluwer.

[29] Cortes-Ciriano, I., Lee, S., Park, W.-Y., Kim, T.-M. & Park, P. J. A molecular portrait of microsatellite instability across multiple cancers. Nature Communications 8, 15180 (2017). URL https://www.nature.com/articles/ncomms15180. Number: 1 Publisher: Nature Publishing Group.

[30] Salipante, S. J., Scroggins, S. M., Hampel, H. L., Turner, E. H. & Pritchard, C. C. Microsatellite instability detection by next generation sequencing. Clinical Chemistry 60, 1192–1199 (2014).

[31] Fujimoto, A., et al. Comprehensive analysis of indels in whole-genome microsatellite regions and microsatellite instability across 21 cancer types. Genome Research 30, 334–346 (2020). URL https://genome.cshlp.org/content/30/3/334. Company: Cold Spring Harbor Laboratory Press Distributor: Cold Spring Harbor Laboratory Press Institution: Cold Spring Harbor Laboratory Press Label: Cold Spring Harbor Laboratory Press Publisher: Cold Spring Harbor Lab.

[32] Verbiest, M. A. et al. Short tandem repeat mutations regulate gene expression in colorectal cancer. Scientific Reports 14, 1–11 (2024). URL https://www.nature.com/articles/s41598-024-53739-0. Number: 1 Publisher: Nature Publishing Group.

[33] Lakshminarasimhan, R. & Liang, G. The Role of DNA Methylation in Cancer. Advances in experimental medicine and biology 945, 151 (2016). URL https://www.ncbi.nlm.nih.gov/pmc/articles/PMC7409375/. Publisher: NIH Public Access.

[34] Ashktorab, H. & Brim, H. DNA Methylation and Colorectal Cancer. Current colorectal cancer reports 10, 425 (2014). URL https://www.ncbi.nlm.nih.gov/pmc/articles/PMC4286876/. Publisher: NIH Public Access.

[35] Zeng, Y., et al. DNA Methylation: An Important Biomarker and Therapeutic Target for Gastric Cancer. Frontiers in Genetics 13 (2022). URL https://www.ncbi.nlm.nih.gov/pmc/articles/PMC8931997/. Publisher: Frontiers Media SA.

[36] Dhar, G. A., Saha, S., Mitra, P. & Chaudhuri, R. N. DNA methylation and regulation of gene expression: Guardian of our health. The Nucleus 64, 259 (2021). URL https://www.ncbi.nlm.nih.gov/pmc/articles/PMC8366481/. Publisher: Nature Publishing Group.

[37] Cunningham, J. M., et al. Hypermethylation of the hMLH1 Promoter in Colon Cancer with Microsatellite Instability. Cancer Research 58, 3455–3460 (1998). URL https://aacrjournals.org/cancerres/article/58/15/3455/504328/Hypermethylation-of-the-hMLH1-Promoter-in-Colon. Publisher: American Association for Cancer Research.

[38] Issa, J.-P. CpG island methylator phenotype in cancer. Nature Reviews Cancer 4, 988–993 (2004). URL https://www.nature.com/articles/nrc1507. Number: 12 Publisher: Nature Publishing Group.

[39] Mousavi, N., Shleizer-Burko, S., Yanicky, R. & Gymrek, M. Profiling the genome-wide landscape of tandem repeat expansions. Nucleic Acids Research 47, e90 (2019). URL https://www.ncbi.nlm.nih.gov/pmc/articles/PMC6735967/.

[40] Mm, H. et al. Frequent MSI mononucleotide markers for diagnosis of hereditary nonpolyposis colorectal cancer. Asian Pacific journal of cancer prevention : APJCP 11 (2010). URL https://pubmed.ncbi.nlm.nih.gov/21133620/. Publisher: Asian Pac J Cancer Prev.

[41] J, W., X, Z., L, J., G, W. & Y, Y. UBR5 Contributes to Colorectal Cancer Progression by Destabilizing the Tumor Suppressor ECRG4. Digestive diseases and sciences 62 (2017). URL https://pubmed.ncbi.nlm.nih.gov/28856538/. Publisher: Dig Dis Sci.

[42] Xie, Z., et al. Significance of the E3 ubiquitin protein UBR5 as an oncogene and a prognostic biomarker in colorectal cancer. Oncotarget 8, 108079 (2017). URL https://www.ncbi.nlm.nih.gov/pmc/articles/PMC5746127/. Publisher: Impact Journals, LLC.

[43] W, D., et al. Inhibition of JAK2/STAT3 signalling induces colorectal cancer cell apoptosis via mitochondrial pathway. Journal of cellular and molecular medicine 16 (2012). URL https://pubmed.ncbi.nlm.nih.gov/22050790/. Publisher: J Cell Mol Med.

[44] Park, S.-Y. et al. The JAK2/STAT3/CCND2 Axis promotes colorectal Cancer stem cell persistence and radioresistance. Journal of Experimental & Clinical Cancer Research 38, 1–18 (2019). URL https://jeccr.biomedcentral.com/articles/10.1186/s13046-019-1405-7. Number: 1 Publisher: BioMed Central.

[45] M, L. & M, G. The emerging role of tandem repeats in complex traits. Nature reviews. Genetics 25 (2024). URL https://pubmed.ncbi.nlm.nih.gov/38714860/. Publisher: Nat Rev Genet.

[46] Mandal, R., et al. Genetic diversity of tumors with mismatch repair deficiency influences anti–PD-1 immunotherapy response. Science 364, 485–491 (2019). URL https://www.science.org/doi/full/10.1126/science.aau0447. Publisher: American Association for the Advancement of Science.

[47] O’Dushlaine, C. T., Edwards, R. J., Park, S. D. & Shields, D. C. Tandem repeat copy-number variation in protein-coding regions of human genes. Genome Biology 6, 1–12 (2005). URL https://link.springer.com/article/10.1186/gb-2005-6-8-r69. Number: 8 Publisher: BioMed Central.

[48] Legendre, M., Pochet, N., Pak, T. & Verstrepen, K. J. Sequence-based estimation of minisatellite and microsatellite repeat variability. Genome Research 17, 1787 (2007). URL https://pmc.ncbi.nlm.nih.gov/articles/PMC2099588/.

[49] Vilar, E. & Gruber, S. B. Microsatellite instability in colorectal cancer—the stable evidence. Nature reviews. Clinical oncology 7, 153–162 (2010). URL https://www.ncbi.nlm.nih.gov/pmc/articles/PMC3427139/.

[50] Siddle, K. J., Goodship, J. A., Keavney, B. & Santibanez-Koref, M. F. Bases adjacent to mononucleotide repeats show an increased single nucleotide polymorphism frequency in the human genome. Bioinformatics 27, 895–898 (2011). URL 10.1093/bioinformatics/btr067. Publisher: Oxford Academic.

[51] Gong, J., et al. PancanQTL: systematic identification of cis-eQTLs and trans-eQTLs in 33 cancer types. Nucleic Acids Research 46, D971–D976 (2018). URL 10.1093/nar/gkx861. Publisher: Oxford Academic.

[52] Jasmine, F. et al. Interaction between Microsatellite Instability (MSI) and Tumor DNA Methylation in the Pathogenesis of Colorectal Carcinoma. Cancers 13, 4956 (2021). URL https://www.ncbi.nlm.nih.gov/pmc/articles/PMC8508563/.

[53] Tomlinson, I., Halford, S., Aaltonen, L., Hawkins, N. & Ward, R. Does MSI-low exist? The Journal of Pathology 197, 6–13 (2002). URL https://pathsocjournals.onlinelibrary.wiley.com/doi/10.1002/path.1071. Publisher: John Wiley & Sons, Ltd.

[54] S, O. & A, G. Molecular classification and correlates in colorectal cancer. The Journal of molecular diagnostics : JMD 10 (2008). URL https://pubmed.ncbi.nlm.nih.gov/18165277/. Publisher: J Mol Diagn.

[55] Mousavi, N., et al. TRTools: a toolkit for genome-wide analysis of tandem repeats. Bioinformatics 37, 731–733 (2021). URL 10.1093/bioinformatics/btaa736. Publisher: Oxford Academic.

[56] Pedregosa, F. et al. Scikit-learn: Machine Learning in Python. Journal of Machine Learning Research 12, 2825–2830 (2011). URL http://jmlr.org/papers/v12/pedregosa11a.html.

[57] Mitra, I., et al. Patterns of de novo tandem repeat mutations and their role in autism. Nature 589, 246–250 (2021). URL https://www.nature.com/articles/s41586-020-03078-7. Number: 7841 Publisher: Nature Publishing Group.

[58] Zhou, H. J., Li, L., Li, Y., Li, W. & Li, J. J. PCA outperforms popular hidden variable inference methods for molecular QTL mapping. Genome Biology 23, 1–17 (2022). URL https://genomebiology.biomedcentral.com/articles/10.1186/s13059-022-02761-4. Number: 1 Publisher: BioMed Central.

[59] J, C.-Z., et al. Comprehensive Analysis of Genetic Ancestry and Its Molecular Correlates in Cancer. Cancer cell 37 (2020). URL https://pubmed.ncbi.nlm.nih.gov/32396860/. Publisher: Cancer Cell.

[60] Seabold, S. & Perktold, J., Walt, S. v. d. & Millman, J. (eds) Statsmodels: Econometric and Statistical Modeling with Python. (eds Walt, S. v. d. & Millman, J.) Proceedings of the 9th Python in Science Conference, 92 – 96 (2010).

[61] Sherman, B. T. et al. DAVID: a web server for functional enrichment analysis and functional annotation of gene lists (2021 update). Nucleic Acids Research 50, W216–W221 (2022). URL 10.1093/nar/gkac194. Publisher: Oxford Academic.

[62] Chen, Y.-a., et al. Discovery of cross-reactive probes and polymorphic CpGs in the Illumina Infinium HumanMethylation450 microarray. Epigenetics (2013). URL https://www.tandfonline.com/doi/abs/10.4161/epi.23470. Publisher: Taylor & Francis.

63. R Core Team. R: A Language and Environment for Statistical Computing (R Foundation for Statistical Computing, Vienna, Austria, 2022). URL https://www.R-project.org/.

[64] Aryee, M. J. et al. Minfi: a flexible and comprehensive Bioconductor package for the analysis of Infinium DNA methylation microarrays. Bioinformatics 30, 1363 (2014). URL https://pmc.ncbi.nlm.nih.gov/articles/PMC4016708/.

[65] Quinlan, A. R. & Hall, I. M. BEDTools: a flexible suite of utilities for comparing genomic features. Bioinformatics 26, 841–842 (2010). URL 10.1093/bioinformatics/btq033. Publisher: Oxford Academic.

[66] Geffen, Y. et al. Pan-cancer analysis of post-translational modifications reveals shared patterns of protein regulation. Cell 186, 3945–3967.e26 (2023). URL https://www.sciencedirect.com/science/article/pii/S009286742300781X.

[67] Roudko, V., et al. Shared Immunogenic Poly-Epitope Frameshift Mutations in Microsatellite Unstable Tumors. Cell 183, 1634–1649.e17 (2020). URL https://www.cell.com/cell/abstract/S0092-8674(20)31463-X. Publisher: Elsevier.

[68] Ballhausen, A., et al. The shared frameshift mutation landscape of microsatellite-unstable cancers suggests immunoediting during tumor evolution. Nature Communications 11, 4740 (2020). URL https://www.nature.com/articles/s41467-020-18514-5. Publisher: Nature Publishing Group.

[69] Os, L., et al. WebSTR: A Population-wide Database of Short Tandem Repeat Variation in Humans. Journal of molecular biology 435 (2023). URL https://pubmed.ncbi.nlm.nih.gov/37678708/. Publisher: J Mol Biol.

